# Mechanism of Curaxin-dependent Nucleosome Unfolding by FACT

**DOI:** 10.1101/2022.05.10.491363

**Authors:** Olesya I. Volokh, Anastasia L. Sivkina, Andrey V. Moiseenko, Anna V. Popinako, Maria G. Karlova, Maria Valieva, Elena Y. Kotova, Mikhail P. Kirpichnikov, Timothy Formosa, Vasily M. Studitsky, Olga S. Sokolova

## Abstract

Human FACT (FACT) is a multifunctional histone chaperone involved in transcription, replication and DNA repair. Curaxins are anticancer compounds that induce FACT-dependent nucleosome unfolding and trapping of FACT in the chromatin of cancer cells (c-trapping) through an unknown molecular mechanism. Here, we analyzed the effects of curaxin CBL0137 on nucleosome unfolding by FACT using spFRET and electron microscopy. By itself, FACT adopted multiple conformations, including a novel, compact, four-domain state in which the previously unresolved NTD of the SPT16 subunit of FACT was localized, apparently stabilizing a compact configuration. Multiple, primarily open conformations of FACT-nucleosome complexes were observed during curaxin-supported nucleosome unfolding. The structures of intermediates suggest “decision points” in the unfolding/folding pathway where FACT can either promote disassembly or assembly of nucleosomes, with the outcome possibly being influenced by additional factors. The data suggest novel mechanisms of nucleosome unfolding by FACT and c-trapping by curaxins.

## Introduction

The eukaryotic genome is organized into nucleosomes *(1,2)* that block DNA accessibility to various sequence-specific DNA-binding proteins. The DNA accessibility is tightly regulated by numerous factors, including ATP-dependent remodelers and ATP-independent histone chaperones *(3-6)*. FACT (facilitates chromatin transcription) is a multifunctional histone chaperone that is involved in both nucleosome assembly and a large-scale, ATP-independent, reversible nucleosome unfolding that increases DNA accessibility to various factors *(7-9)*. These activities of FACT contribute to various nuclear processes including transcription, replication and repair *(6,9,10)*.

FACT is a conserved protein complex *(6)* that consists of two subunits: SPT16 (Suppressor of Ty 16) and SSRP1 (Structure Specific Recognition Protein 1) in human and plants; Pob3 (Polymerase One Binding protein 3) replaces SSRP1 in yeasts. Human SPT16 and SSRP1 subunits consist of four and five structurally different regions, respectively (Fig. 1A), implicated in binding to both nucleosomal DNA and to different core histones (see *(3-6)* for review).

**Fig. 1.**
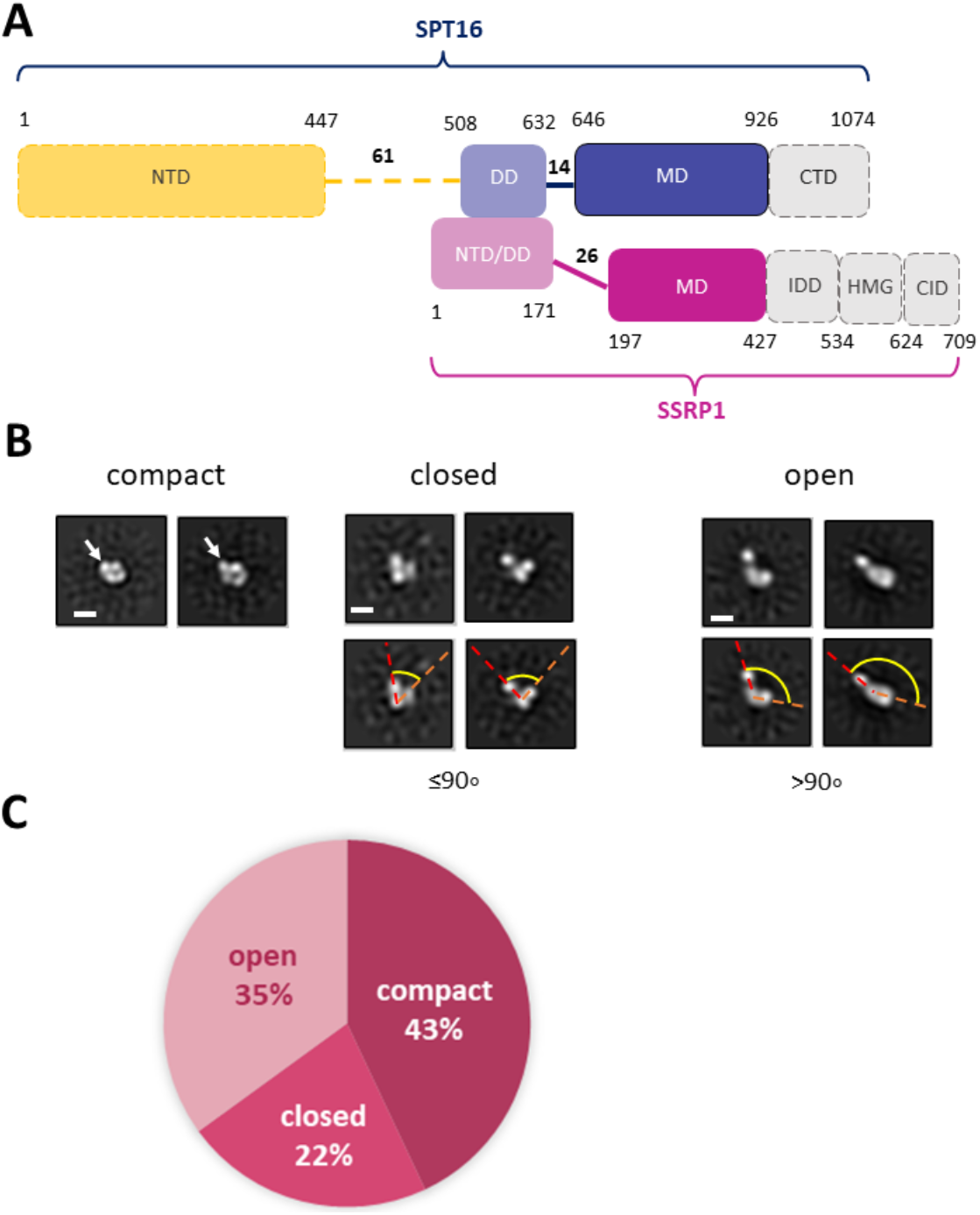
Human FACT is a flexible protein complex. (**A**) Human FACT domain architecture. Domains with dashed boundaries are expected to be disordered; indeed, domains in gray are not detectable by TEM. Domain abbreviations: NTD – N-terminal, DD – dimerization, MD – middle, CTD – C-terminal, HMG – high mobility group box, IDD – intrinsically disordered, CID - CTD-Interacting domain, (**B**) Characteristic 2D class averages of FACT in compact, closed and open conformations: the arrow indicates the domain that is detectable only in the compact conformation, (**C**) Distribution (%) of FACT in different conformations.

FACT inhibition negatively affects a number of the critical p53-, NF-kB- and HSF1-dependent metabolic pathways involved in cancer development and leads to the death of the cancer cells *(11)*. Human FACT (FACT) is overexpressed in various types of cancer and thus is a promising target for anti-cancer drugs *(11-13)*. In particular, members of the class of DNA intercalators called curaxins have strong anticancer activity (*11,14*), induce chromatin trapping (c-trapping) of FACT and strongly inhibit normal human FACT activities *in vivo (11)*. C-trapping of FACT involves formation of Z-DNA *in cellulo (15)* and curaxin-dependent nucleosome unfolding accompanied by tight binding of FACT to the unfolded nucleosomes *in vitro (16)*. The anticancer activity of curaxins is highly dependent on c-trapping of FACT *(11,15,16)*.

Recently, using transmission electron microscopy we have described a number of topologically different structures of yeast FACT(yFACT)-nucleosome complexes formed during nucleosome unfolding by yFACT and Nhp6 protein *(7)*. Our data suggested that recently determined high resolution structures of human FACT bound to subnucleosomal complexes *(17,18)* are structurally similar to the early intermediates formed during nucleosome unfolding. It was also shown that in the unfolded complexes yFACT is engaged in multiple interactions, both with nucleosomal DNA and core histones *(7)*, raising the possibility that a similar mechanism could be involved in curaxin-dependent nucleosome unfolding by human FACT.

Here spFRET microscopy, single particle transmission electron microscopy (TEM) and molecular modeling (MM) revealed high conformational flexibility of both FACT and FACT-nucleosome complexes formed in the presence of the clinically relevant curaxin CBL0137. The structures of the intermediates formed during the FACT/curaxin-dependent nucleosome unfolding illuminate the mechanism of this process and the mechanism of c-trapping of FACT by curaxins.

## Results

### Human FACT is a highly flexible protein complex

The structure of human FACT (FACT, Fig. 1A) was studied using transmission electron microscopy (TEM). To ensure correct alignment of the particles, they were classified in RELION2.1 (Table. S1, Fig. S1). TEM revealed three distinct conformations of FACT: compact, closed and open (Fig. 1B). The closed conformation includes three closely positioned densities that dissociate from one another in the open conformation (Fig. 1B). These complexes are structurally similar to those detected previously during TEM studies of yeast FACT *(7)*. Here TEM also revealed a novel, abundant, compact conformation of FACT that consists of four globular densities arranged in a compact diamond-like shape (Fig. 1B) that was not detected previously with yeast FACT *(7)*. The three conformations were present at ratios of approximately 2:1:1.6 (compact:closed:open) (Fig. 1C).

Based on the previous identification of the components of yFACT *(7)*, the three densities present in all conformations of FACT were tentatively identified as SSRP1-NTD/DD-SPT16-DD, SSRP1-MD and SPT16-MD (Fig. 1A). Accordingly, the 2D projections of those densities are ∼4-5 nm in diameter, as expected for these structures with molecular masses of ∼30-40 kDa. The fourth electron density, detectable only in the compact conformation, is somewhat larger (∼5-6 nm in diameter, Fig. 1B), and is therefore likely to be the Spt16-NTD domain, which at ∼50 kDa is the largest domain of FACT (*19*). This domain was not detected in previous EM studies of FACT and FACT-nucleosome complexes (*7,17*), presumably indicating larger conformational flexibility of this region in the structures detected.

To further evaluate whether the additional density is indeed the NTD of SPT16, we analyzed a truncated version of FACT lacking this domain (SPT16ΔNTD) by TEM. With this construct, only closed (48%) and open (52%) conformations were identified, with no 2D class corresponding to the compact 4-lobed structure being observed (Fig. S2A). The data are consistent with the proposal that the largest electron density detected in the compact conformation of FACT (Fig. 1B) is indeed the SPT16-NTD domain.

### Domain identification in 3D structures of FACT

To identify FACT domains more directly and to ensure that the observed conformational states do not simply reflect different orientations of the same configuration, 3D maps of FACT in the compact and open conformations, and in the closed conformation of FACT containing SPT16ΔNTD that produced more homogeneous and better resolved complexes (Fig. 2A-C). The resolutions of the reconstructions were moderate (21Å for compact, 34Å for closed and 31Å for open conformations, respectively), reflecting high flexibility of the FACT molecule. The linear dimensions of FACT are 12±0.4 × 8.6±0.7 nm for the compact, 9.1±0.4 × 5.4±0.5 nm for the closed and 15±1.9 × 5.5±0.6 nm for the open conformations, respectively.

**Fig. 2.**
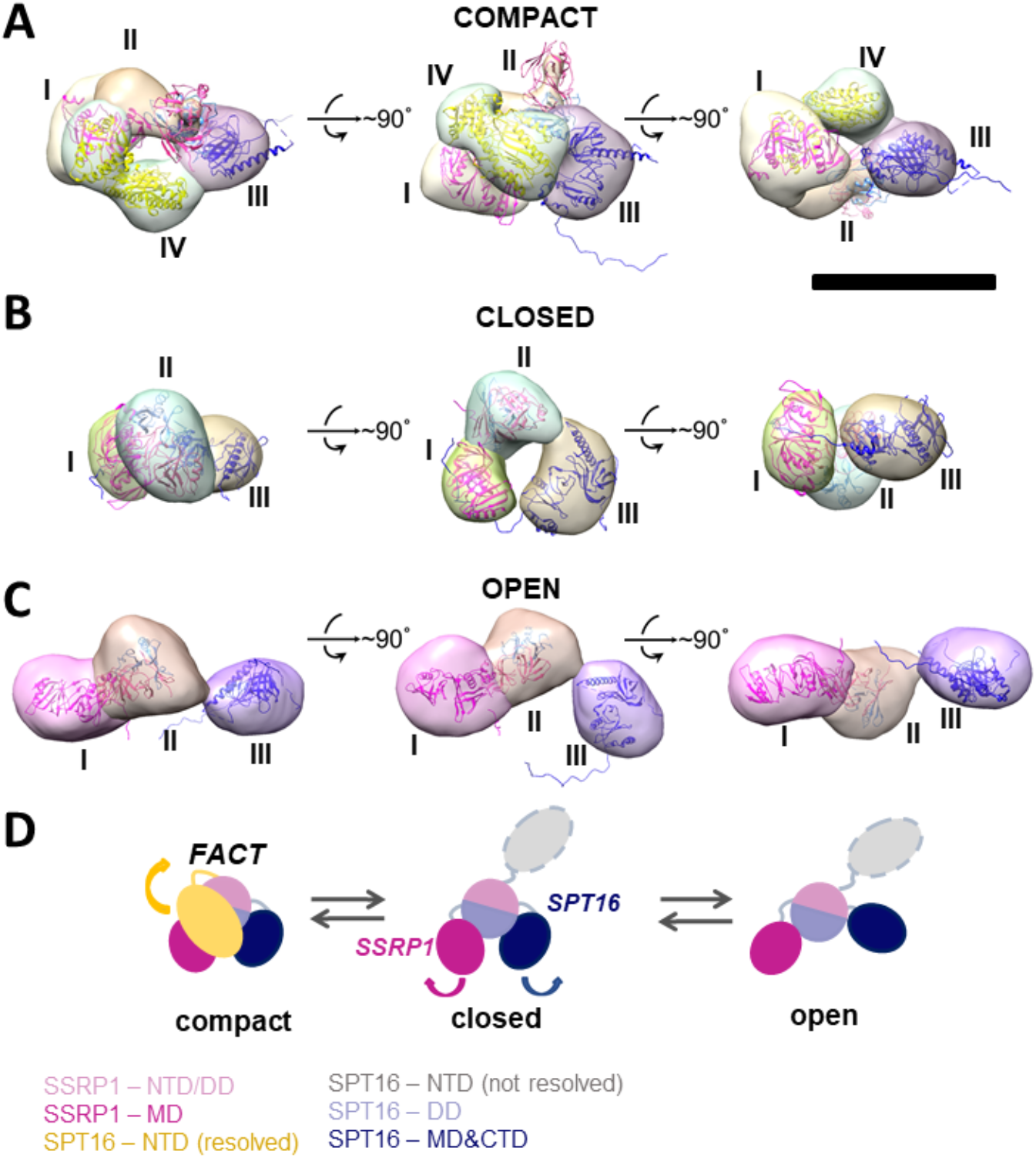
Three-dimensional structures of FACT in compact, closed and open conformations. (**A**)-(**C**). 3D structures of FACT in the compact (**A**), closed (**B**) and open (**C**) conformations. In the compact complex, the crystal structures of the domains were docked into corresponding EM densities I-IV with correlation coefficients of 0.92; 0.91; 0.89; 0.93, respectively. Bar – 10 nm. d. The proposed pathway of FACT unfolding. The color code for the domains of FACT is shown at the bottom of the figure.

To localize the domains in the electron densities of FACT, rigid fitting was performed using the available crystal structures of FACT domains (Fig. 2). Based on previous EM studies (*7,17*), we assumed that the middle density is SSRP1-NTD/DD-SPT16-DD (domain II), while the SSRP1-MD (domain I) and SPT16-MD (domain III) flank it on either side (Fig. 2). This structural assignment results in a good fit of all domain structures into the electron densities of the best resolved compact conformation of FACT (Fig. 2A). For the closed and open FACT conformations the MD domains of SSRP1 and SPT16 were positioned into the densities I and III based on the length of the linkers connecting the domains. Because the linker connecting NTD/DD and MD domains of SSRP1 is longer than the one connecting NTD/DD and MD of SPT16 (Fig. 1A), SSRP1-MD (domain I) is likely to be connected with the NTD/DD through a less extensive electron density than SPT16-MD (Fig. 2B, C). Crystal structures were automatically fitted into corresponding domains with correlation coefficients >0.89.

To localize the SPT16-NTD domain, the 3D map of compact conformation of full-length FACT (Fig. S2B) was first aligned with the map of closed conformation of the FACT SPT16ΔNTD mutant (Fig. S2C). The difference map revealed an additional density in FACT in comparison with the mutant version of the complex (shown in magenta mesh in Fig. S2C). Rigid fitting of the crystal structure of SPT16-NTD (pdb ID 5e5b *(20)*) into this density yielded good correspondence of the structures, with correlation coefficient 0.92 (Fig. S2C).

To evaluate possible driving forces allowing formation of the compact FACT conformation (Fig. 2A) the interacting surfaces of all subunits were analyzed using flexible molecular docking of SPT16-NTD to other resolved domains in the compact conformation of FACT with the correlation coefficient 0.89 using HADDOCK (Fig. S3). In the resulting model, three domains of FACT (SSRP1-NTD/DD, SPT16-DD and SPT16-NTD) are tethered together through hydrophobic interactions between the subunits with the buried surface contact area of ∼2004 Å^2^ (Fig. S3). Thus, the hydrophobic interactions that connect different domains in the compact FACT complex are quite strong and are potentially able to stabilize FACT in a compact conformation in solution. The hydrophobic interactions are supplemented by a dense network of hydrogen bonds between the domains (Table. S2).

Since no open complexes with four densities were observed, the data suggest that SPT16-NTD domain “locks” the other domains of FACT in the compact conformation (Fig. 2A), primarily through hydrophobic interactions supplemented by multiple hydrogen bonds (Fig. S3). When the NTD is displaced, the remaining domains form a more flexible structure that is in equilibrium between the open and closed states (Fig. 2D).

The SPT16-NTD domain is not detectable in the other closed conformations or in any of the open forms, most likely because once it dissociates from the remaining FACT complex it becomes more mobile and therefore “invisible” in the 2D and 3D class averages; indeed, it was not detected in previous structural studies of FACT-nucleosome complexes *(17)*. Alternatively, separation of any density from the compact complex could induce separation and mobilization of the SPT16-NTD domain of FACT. This possibility is unlikely because it predicts that all complexes with three densities would be in an open state, but we also observed closed three-density complexes (Fig. 1B).

In summary, TEM revealed that FACT is a mixture of three conformations: compact, closed and open. Four or three distinct densities are visible in the compact and closed/open conformations, respectively. The three densities were identified as SSRP1-MD (domain I), SSRP1-NTD/DD-SPT16-DD (domain II) and SPT16-MD (domain III); the fourth domain is SPT16-NTD. The arrangement of the densities in the complexes suggests that SPT16-NTD domain “locks” the other domains of FACT in the compact conformation.

### Nucleosome unfolding by FACT in the presence of curaxin CBL0137

The interaction of FACT with nucleosomes was studied using mononucleosomes assembled on the 603 Widom nucleosome positioning sequence *(21)*. Nucleosomal DNA contained a single pair of Cy3 and Cy5 fluorophores in positions 35 and 112 bp from the nucleosomal entry/exit boundary, allowing fluorescence resonance energy transfer (FRET) between the fluorophores and detection of the conformation changes in nucleosomal DNA upon interaction with FACT and curaxins (*9,16*). Single particle FRET (spFRET) from the nucleosomes was measured in the absence and presence of curaxin CBL0137, FACT and competitor DNA (Fig. 3A).

**Fig. 3.**
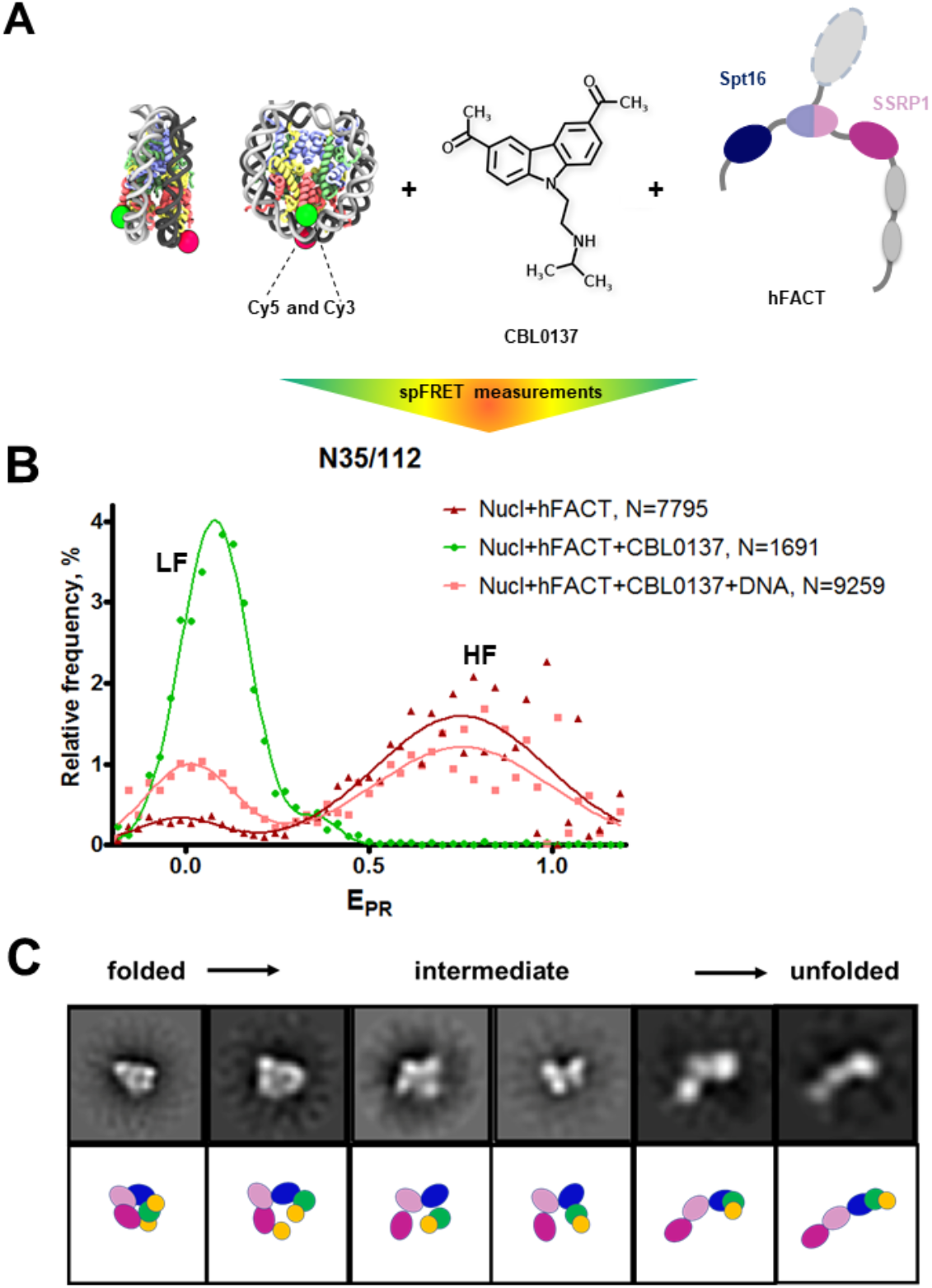
FACT and curaxin CBL0137 work synergistically and induce a large-scale nucleosome unfolding. (**A**) Experimental approach for analysis of nucleosome unfolding using spFRET. Mononucleosomes contained a single pair of Cy3 and Cy5 dyes in the nucleosomal DNA (shown as green and red circles, respectively), (**B**) Nucleosome unfolding by FACT/CBL0137 under conditions used for electron microscopy: analysis by spFRET microscopy. Frequency distributions of FRET efficiencies (E_PR_) in the presence of FACT and/or CBL0137 and competitor DNA. Gaussian peaks having lower and higher FRET efficiencies are indicated (LF and HF, respectively), (**C**) Nucleosome unwrapping by FACT in the presence of CBL0137. Top: Characteristic 2D class averages of FACT-nucleosome complexes in the presence of CBL0137. The complexes are arranged to show the proposed sequence of events during nucleosome unfolding by FACT. Bar – 10 nm. Bottom: The observed electron densities are shown in different colors to illustrate the conformational transitions.

As expected, no changes in nucleosome structure were detected in the presence of FACT alone and only minor increase of the height of the low-FRET peak and corresponding decrease of the high-FRET peak were detected in the presence of CBL0137 only (Fig. S4). In contrast, FACT and CBL0137 added to the nucleosomes together induced a profound transition from high to low FRET, reflecting a dramatic uncoiling of nucleosomal DNA (Figs. 3B and S4) (*9,16*). These changes in the structure of nucleosomal DNA were largely reversed by subsequent addition of an excess of competitor DNA that removes FACT from the complex. Thus, FACT induces a large-scale, reversible nucleosome unfolding in the presence of curaxin (*16,22*).

To directly visualize the process of nucleosome unfolding by FACT in the presence of CBL0137, the complexes of FACT with nucleosomes were formed in the presence of CBL0137, characterized by spFRET microscopy immediately before EM (Fig. 3B and Fig. 3C), applied to the EM grid, negatively stained and studied using TEM. Single particle images were collected using a neural network in EMAN2.3 *(23)* and subjected to 2D-classification in RELION2.1 (Figs. S5 and S6).

In the sample that contains FACT and nucleosomes in the absence of curaxin the following class-average complexes were detected after 2D classification (Fig. S5): i) nucleosomes (an excess of nucleosomes was added to minimize the presence of nucleosome-free FACT), ii) nucleosome-free FACT present in the open, compact and closed conformations, and iii) folded FACT-nucleosome complexes.

Adding curaxins to the FACT-nucleosome complex resulted in formation of several novel conformations of the complex (Figs. 3C and S6), which are likely to represent intermediates formed during stepwise nucleosome unfolding. Multiple intermediates between the initial folded and fully unfolded complexes were identified (Figs. 3C and S7); the length of the intermediates spanned the range from 17.3±2.3 nm to 21.6±2.5 nm.

Importantly, the 2D projections of the folded FACT-nucleosome complexes closely resembled those obtained by Liu. et al *(17)* (Fig. 4A), although entirely different strategies for assembly of the complexes were used. This observation allowed reconstruction of the 3D structure of the compact FACT-nucleosome complexes using RELION 3.0; the 3D model was built using 19,074 particles with a final resolution of 22Å (Fig. 4B). The previously determined atomic structure of the folded FACT-nucleosome complex *(17)* was fitted in the observed 3D electron density with the correlation coefficient of 0.92, indicating similar structures of the complexes.

**Fig. 4.**
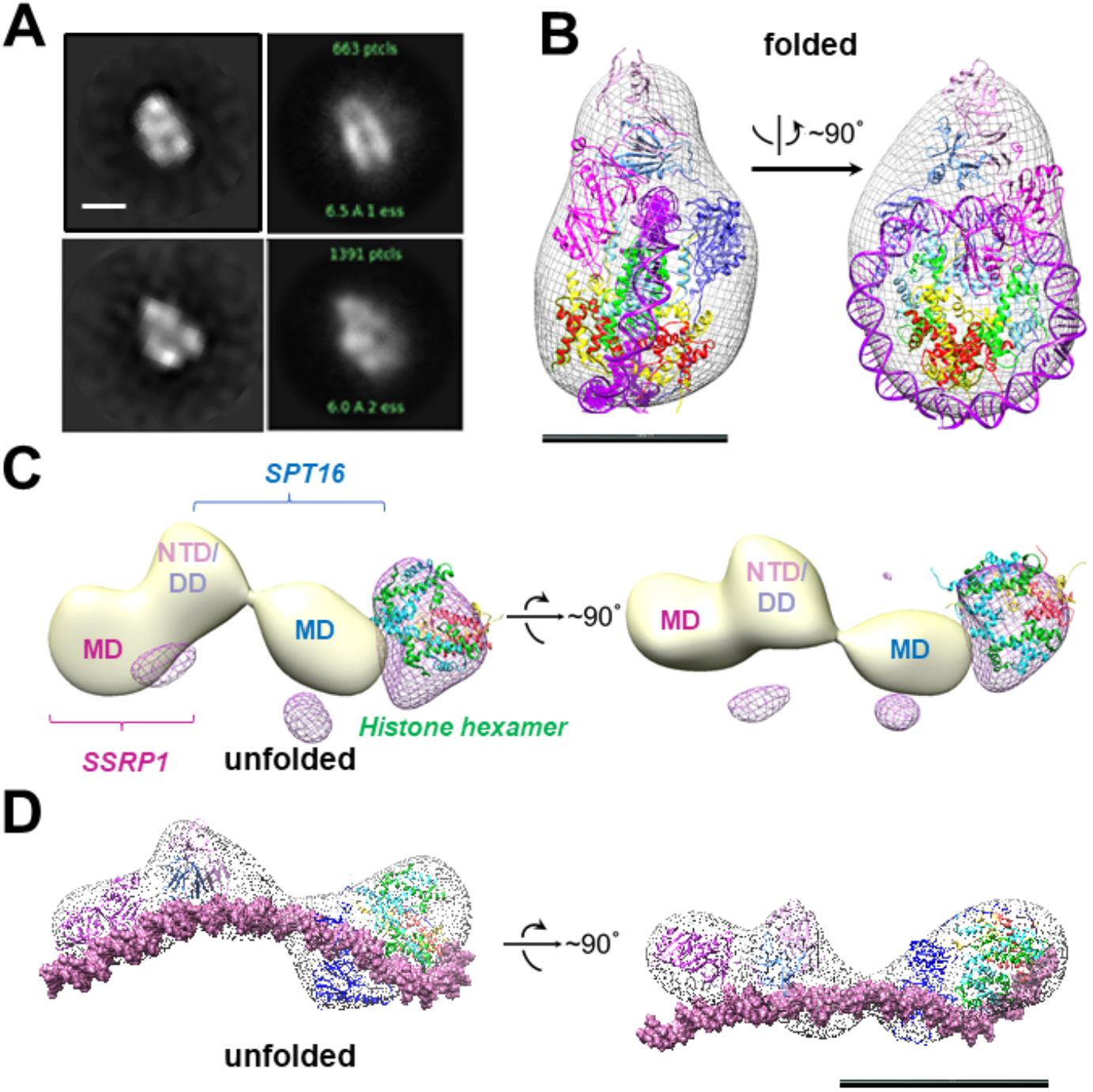
Structures of folded and unfolded FACT-nucleosome complexes formed in the presence of curaxin CBL0137. (**A**) Comparison of similar 2D class-averages of folded FACT-nucleosome complexes from the present study (left) and from Liu et al. (*17*) (right). Scale bar – 10 nm, (**B**) A model of the compact FACT-nucleosome complex with crystal structures of SPT16-DD, SSRP1-DD, SSRP1-MD, SPT16-NTD/MD/CTD (PDB 6UPL) fitted into the electron densities of the complex with the correlation coefficient 0.92, (**C**) Difference map between unfolded FACT-nucleosome complex and FACT in the open state (light yellow); the resulting differential density is shown in magenta mesh. Histone hexamers (H2A/H2B dimer and H3/H4 tetramer derived from 6UPK PDB) were fitted into the differential density with correlation coefficient of 0.88, (**D**) A model of the unfolded FACT-nucleosome complex. DNA is positioned based on the assumption that both SPT16-DD/MD and SSRP1-DD/MD domains maintain their interactions with DNA previously observed in the compact complex (*17*).

The images of the unfolded complexes were extracted from the dataset and used for 3D reconstruction of the unfolded complex in RELION2.1 (Fig. 4C). The reconstruction has a clear four-density structure, with three densities similar in size to corresponding densities of FACT in the open conformation (compare Figs. 2C and 4C), and the additional fourth domain (Fig. S8). The fourth domain can accommodate the H3/H4 tetramer and possibly one H2A/H2B dimer (linear dimensions are ∼10×5 nm, Fig. 4C). Assuming that DNA in the unfolded complex is nearly linear, FACT and a histone tetramer are bound to ∼80-bp DNA region (Fig. 4D). The second H2A/H2B dimer could remain in contact with SSRP1-MD domain, stabilized by SSRP1-CID region and flexibly linked to nucleosomal DNA *(24)*, preventing it from being resolved in the open complex. Other FACT domains (SPT16 CTD, SSRP1 IDD&HMG&CID) are also unlikely to be ordered sufficiently to be resolved *(17)*.

In summary, binding of FACT to the nucleosome in the presence of curaxin CBL0137 induced a dramatic unfolding of nucleosomal DNA that was accompanied by formation of a multi-density complex containing core histones and both subunits of FACT. The complex is a mixture of intermediates that contain nucleosomes unfolded to different degrees. The most folded complex is structurally similar to the FACT-nucleosome complex characterized previously *(17)*; the similarity allowed assignment of electron densities in the folded complex to various FACT domains and core histones. Subsequent analysis of the unfolded intermediates suggests a pathway of progressive curaxin-dependent nucleosome unfolding by FACT.

### Mechanism of FACT/curaxin-dependent nucleosome unfolding

The data described above suggest the following scenario for nucleosome unfolding by FACT in presence of curaxin (Fig. 5). Nucleosome-free FACT a mixture of compact, closed and open states (Fig. 1B). In the compact conformation of the complex, the C-terminal DNA-binding regions of both subunits of FACT could interact with other domains of FACT *(7)* and the DNA-binding domains on the SPT16 and SSRP1 subunits are likely hidden and not available for interaction with a nucleosome.

**Fig. 5.**
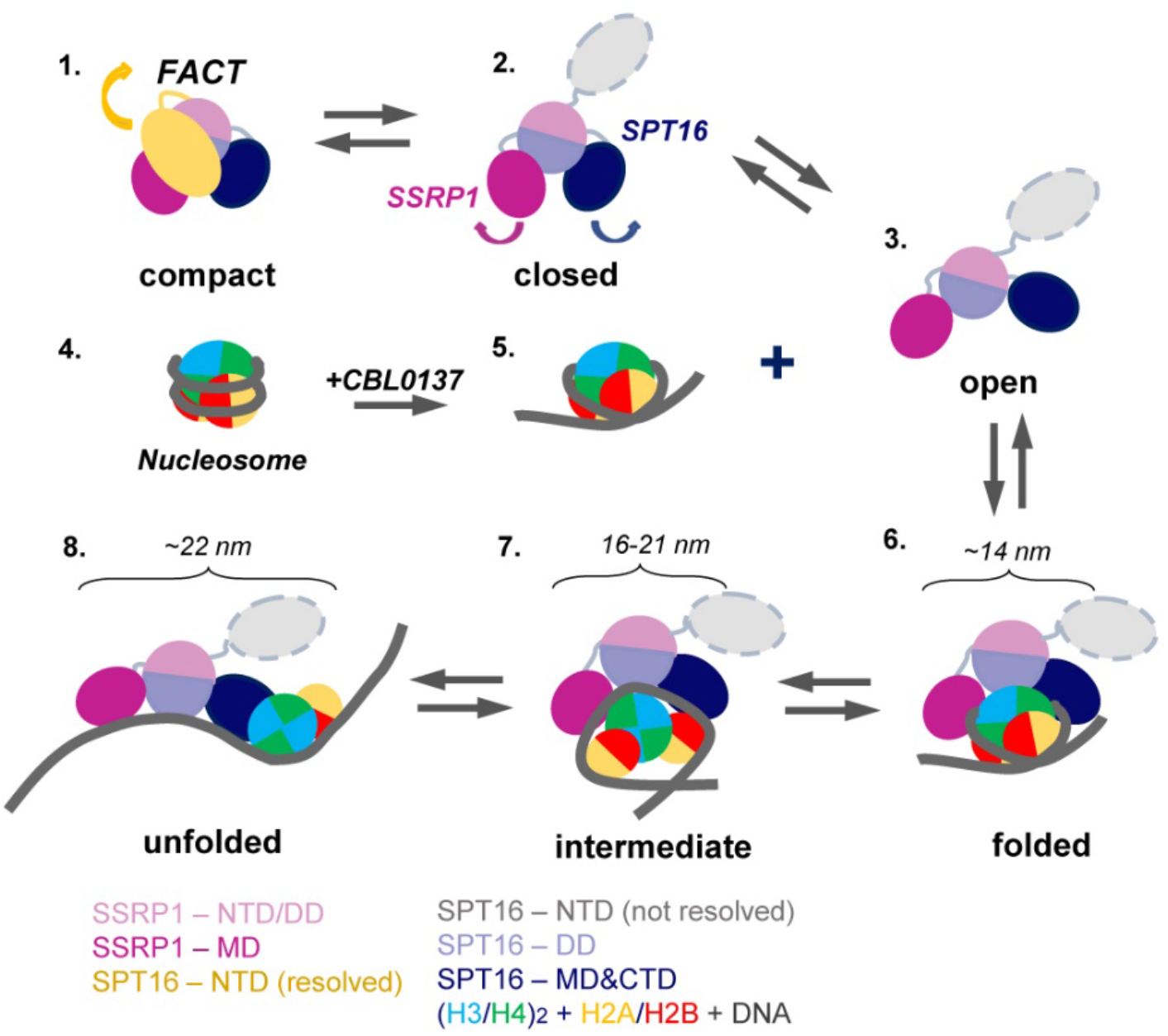
Model of nucleosome unfolding by FACT in the presence of curaxin CBL0137. FACT is a mixture of compact, closed and open states (intermediates 1, 2, and 3). As nucleosomal DNA is partially uncoiled from the histone octamer in the presence of CBL0137 (intermediates 4 and 5), FACT-binding sites on the surface of H2A/H2B dimers become available for interaction with FACT, and the folded complex is formed (intermediate 6). As a result, FACT induces further nucleosome unfolding (intermediates 7 and 8). The linear dimensions of the intermediates 6 to 7 are indicated. The color code is shown at the bottom.

The comparison of 3D structures of FACT (Fig. 2B, C) suggests that opening of the complex likely occurs through a concerted movement of the four domains. First, SPT16-NTD moves away from the complex and thus probably “unlocks” the mobilities of the other domains (Fig. 5, intermediates 1 and 2); then other domains of FACT move away from each other, forming the open complex (Fig. 5, intermediate 3). In the open conformation of FACT the DNA-binding sites on the dimerization and middle domains of SPT16 and SSRP1 subunits, respectively *(17)*, as well as the C-terminal DNA-binding regions of both subunits of FACT *(7)* become available for interaction with nucleosomes.

However, FACT interacts weakly with intact nucleosomes; the DNA at the entry/exit sites has to be partially displaced from the histone octamer to expose binding sites for FACT, which was accomplished by removing this DNA in the cryo-EM structure *(17)* or by adding high levels of the HMBG factor Nhp6 *(7)*. Our results show that curaxin CBL0137 provides this activity as well (Fig. 5, intermediates 4 and 5). This DNA intercalator *(15)* binds to and induces partial displacement of the nucleosomal DNA from the octamer (Fig. 3B and *(16)*), exposing the FACT-binding surfaces on H2A-H2B dimers. FACT binds to the destabilized nucleosome and the folded complex is formed (Fig. 5, intermediate 6); the complex is structurally similar with the FACT-nucleosome complex described previously *(17)*.

The initial binding of FACT to the nucleosome triggers a progressive sequence of events leading to formation of the intermediate and unfolded complexes (Fig. 5, intermediates 7 and 8) containing nearly completely uncoiled nucleosomal DNA. Since FACT-dependent nucleosome unfolding is an ATP-independent process (*7,16*), it most likely occurs through a set of intermediates (Fig. 3C) having similar free energies that are reversibly interconverted. For each pair of the intermediates the equilibrium can be easily shifted in either direction by engaging additional protein-protein and/or DNA-protein interactions *(7)*, or through partial uncoiling of nucleosomal DNA from the octamer by curaxins.

The unfolded complex is stabilized by multiple interactions of different FACT domains with both nucleosomal DNA and core histones (Fig. 4D). Since each of these interactions is relatively weak, nucleosome unfolding is a partially reversible process: thus, intact nucleosomes can be largely recovered in the presence of competitor DNA (Fig. 3B) that presumably binds and outcompetes the curaxin from the nucleosomal DNA. Upon the removal of the curaxin nucleosomal DNA re-binds to core histones and FACT dissociates from the complexes.

## Discussion

Our structural analysis of the process of curaxin-dependent nucleosome unfolding by FACT using spFRET and TEM revealed that in the absence of nucleosomes FACT is a flexible complex that exists in compact, closed and open conformations present at the ratio of 43:22:35, respectively (Fig. 1). Four or three distinct densities are visible in the compact and closed/open conformations, respectively; molecular modeling allowed assignment of these electron densities to FACT domains (Fig. 2). The arrangement of the densities in the complexes suggests that SPT16-NTD domain “locks” the other resolved domains of FACT in the compact conformation (Fig. 2D). While FACT alone binds weakly to a nucleosome, multiple structurally different FACT-nucleosome complexes (folded, intermediate and unfolded) are formed in the presence of curaxin CBL0137 (Fig. 3). Electron densities in the folded complex were assigned to several FACT domains and core histones (Fig. 4A, B). Molecular modeling suggests that the unfolded complex contains nearly linear DNA, core histones and both subunits of FACT (Fig. 4C, D). Subsequent analysis of the unfolded intermediates suggests a pathway of progressive curaxin-dependent nucleosome unfolding by FACT (Fig. 5).

Similar distributions between the open and closed conformations for yeast (*7*) and human FACT in the absence of other factors (∼35:65, Fig. S9) highlight the overall structural similarity of these factors. At the same time, the pathways of nucleosome unfolding by yeast FACT in the presence of the DNA-binding protein Nhp6 (*7*) and by human FACT in the presence of curaxin are different. In both cases, complete nucleosome unfolding requires the presence of all participating factors. However, the curaxin interacts with DNA and therefore the observed increase in the presence of the open FACT complexes after the nucleosome unfolding from 35 to 44% (Fig. S9A) likely occurs only due to nucleosome destabilization by curaxin (Fig. S4). In contrast, Nhp6 protein interacts both with FACT and with nucleosomes (*7*), and therefore induces an increase in the fraction of the open forms of FACT both in the absence and in the presence of nucleosomes (36 to 51% and to 55%, respectively, Fig. S9B). Accordingly, the overall efficiencies of nucleosome unfolding by FACT with curaxin and with Nhp6 protein are different (44 vs. 55%, respectively).

Comparison of the curaxin- and Nhp6-dependent pathways also suggests that partial uncoiling of nucleosomal DNA from the histone octamer is a necessary pre-requisite for nucleosome unfolding by FACT. Partial DNA uncoiling that must occur during transcription and replication exposes FACT-binding sites on the octamer and provides a target for FACT binding. Indeed, FACT is associated with transcribed genes and the replication fork (*25-28*); since destabilized nucleosomes have exposed binding sites for FACT, nucleosome unfolding likely occurs during these processes (Fig. S10A). It has been proposed that nucleosome unfolding could facilitate nucleosome survival during transcription (*29*); although there is no direct evidence for unfolding of these nucleosomes, FACT facilitates nucleosome survival during transcription *in vitro* (*28*). Therefore, FACT could possibly induce nucleosome unfolding and this could support nucleosome survival (Fig. S10A).

At the same time, FACT is known for its ability to both destabilize and assemble nucleosomes. Formation of energetically similar intermediates during FACT-dependent nucleosome unfolding (Fig. 5, intermediates 6-8) can explain this apparent contradiction in properties. Indeed, the equilibrium between the intermediates can be easily shifted in either direction in processes involving relatively low energy cost, such as formation of several additional DNA-protein interactions by Nhp6 protein (*7*) or curaxin intercalation into nucleosomal DNA (this work). These factors induce FACT-dependent nucleosome unfolding; an example of the process developing in the opposite direction is reversal of FACT-dependent nucleosome unfolding in the presence of competitor DNA (Fig. 3B). None of these factors (Nhp6, curaxin or competitor DNA) strongly affects intact nucleosomes; all of them work highly synergistically with FACT. In all cases FACT creates an intermediate that serves as a “decision point” allowing either nucleosome unfolding or recovery after an additional interaction of the factors with DNA or a protein.

Curaxins reduce the growth of cancer cells by inducing the trapping of FACT within bulk chromatin (c-trapping) (*11,16*). The trapping most likely occurs because human FACT binds weakly to intact nucleosomes (Fig. S7) but has higher affinity for partially unwound nucleosomes. Curaxin-induced binding of FACT to bulk nucleosomes (Fig. S7) most likely explains the observed re-distribution of FACT from transcribed genes to bulk chromatin (*16*). Importantly, the dramatic curaxin- and FACT-induced nucleosome unfolding (Fig. 3) likely changes the global arrangement of topologically closed chromatin loops, introducing unconstrained negative DNA supercoiling (*22*). The changes in chromatin topology likely contribute to the observed genome-wide formation of Z-DNA in cancer cells in the presence of curaxins (*15,30*) (Fig. S10B).

A limitation of our electron microscopy approach is the use of negative stain that was used because the analyzed structures are highly flexible *(7)*; therefore the resolution is limited by the contraster grain size. However, since the critical structures obtained in our studies are very similar with the structures obtained using more preservative electron cryo-microscopy (Fig. 4A) the molecular details of the mechanism can be derived with confidence.

In summary, FACT is a remarkably flexible protein complex; its flexibility is an important factor during FACT-dependent nucleosome unfolding where “decision point” intermediates are formed. The equilibrium between the intermediates can be shifted in either direction with relatively low energy cost. Similar intermediates are likely formed during transcription and replication in the cell nuclei; their formation allows FACT to either destabilize or assemble nucleosomes, depending on the presence of additional factors. In particular, curaxins induce reversible FACT-dependent nucleosome unfolding, likely through intercalation into nucleosomal DNA and destabilization of nucleosomes, leading to FACT trapping in bulk chromatin in cancer cells.

## Materials and Methods

### Experimental Design

The objectives of the study were to determine the molecular mechanism of curaxin-dependent nucleosome unfolding by FACT. The experimental design includes use of highly purified components (core nucleosomes, curaxin CBL0137 and FACT) and experimental approaches allowing detailed analysis of highly flexible complexes (electron microscopy and spFRET).

### Protein purification

-H1 chicken erythrocyte chromatin and chicken nucleosome cores were purified as described (*31*). Recombinant FACT subunits were co-expressed in insect Sf9 cells as described (*25*) and FACT was purified as described (*28*). Recombinant SPT16/SSRP1ΔNTD complex was expressed as a dimer in *E*.*coli* and purified as described (*9*) .

### Nucleosome assembly and purification

A plasmid containing the modified 603-42 nucleosome positioning sequence (*32*) was used to obtain the nucleosomal DNA template by PCR with the following fluorescently labeled primers:

#### Forward primer

5′–CCCGGTTCGCGCTCCCTCCTTCCGTGTGTTGTCGT*CTCT-3’ (where T* - is a nucleotide labeled with Cy5);

#### Reverse primer

5′– ACCCCAGGGACTTGAAGTAATAAGGACGGAGGGCCT#CTTTCAACATCGAT-3’ (where T# - is a nucleotide labeled with Cy3).

The 147-bp N35/112 DNA fragments were purified using Evrogen Cleanup Standart kit (Evrogen, Russia).

Nucleosomes were assembled by octamer transfer from -H1 chromatin to DNA templates after dialysis from 1M NaCl to 0.01M NaCl as described (*31,33*). For gel shift analysis of FACT binding and spFRET analysis nucleosomes were purified by PAGE under non-denaturing conditions as described (*9,34*).

For spFRET experiments N35/112 nucleosomes were gel purified and analysed at a concentration of 0.5-1 nM after incubation in the presence of FACT (0.1 µM) and/or CBL0137 (2 µM) in the buffer containing 20mM Tris-HCl pH7.9, 150 mM KCl for 5 min at 25°C.

#### spFRET experiments

spFRET measurements in solution and analysis were performed as described (*9,35*). The proximity ratio E_PR_ was calculated as

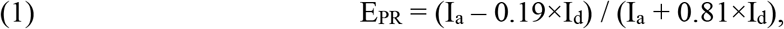

where I_a_ and I_d_ are Cy5 and Cy3 fluorescence intensities corrected for background. Factors 0.19 and 0.81 were introduced to correct for the contribution of Cy3 fluorescence in the Cy5 detection channel (spectral cross-talk) (*35*).

Proximity ratios E_PR_ were calculated using 800-8000 signals from single nucleosomes for each measured sample and plotted as a relative frequency distribution. Each plot was fitted with a sum of two Gaussians to describe two conformational states of nucleosomes. The fractions of nucleosomes in different states were estimated as the areas under the corresponding Gaussian peaks normalized to the total area of a plot by using LabSpec program. Reproducibility of the results was verified in at least three independent experiments.

### Electrophoresis mobility shift assay (EMSA)

All EMSA experiments were performed using the native 4% PAGE in 0.5×TBE buffer (40 mM Tris-Cl, pH 8.3, 45 mM boric acid, 1 mM EDTA) under 4°C conditions.

### Preparation of samples for electron microscopy

SPT16/SSRP1 and SPT16/SSRP1ΔNTD complexes were prepared in previously characterized buffer containing 17 mM HEPES pH 7.6, 2 mM Tris-HCl pH 7.5, 0.8 mM Na3EDTA, 0.11 mM 2-mercaptoethanol, 11 mM NaCl, 1.1% glycerin, 12% sucrose (*7,9,28*) at concentration of 0.05 µM.

Complexes of FACT with the nucleosome were formed in the presence of 0.1 µM FACT, 0.1 µM core chicken nucleosomes, 2 µM CBL0137 and 0.5 nM fluorescently labeled core nucleosomes N35/112 on ice for 42 h. Since complexes of human FACT with nucleosomes are not stable in a native gel but stable in solution (*16*), the complexes were analyzed using spFRET and TEM. Part of the sample was used to evaluate reversibility of nucleosome reorganization by adding salmon sperm DNA to final concentration 0.65 µg/µl for 0.5 h on ice, followed by spFRET-microscopy. The remaining sample was used for TEM.

### Transmission electron microscopy and image analysis

Protein samples and complexes were applied to the carbon-coated glow-discharged in Emitech K100X device (Emitech Ltd., UK) copper grid (Ted Pella, USA) immediately after preparation, subjected to glow-discharge using Emitech K100X device (Emitech Ltd., UK), stained for 30 sec with 1% uranyl acetate, and air dried. Grids were studied in JEOL 2100 TEM (JEOL) microscope operated at 200 kV at low-dose conditions. Micrographs were captured by the Gatan Ultrascan camera with magnification x25,000, no tilt, with 4.1 Å pixel size using SerialEM software (*36,37*).

Single particle images of FACT, complexes of FACT with the nucleosome and complexes of FACT with the nucleosome formed in the presence of CBL0137 were collected from the micrographs using a neural network provided by EMAN2.3 software. Single particles coordinates collected by the neural network were imported in RELION2.1 software; all further 2D-processing, analysis and CTF-correction were performed using RELION2.1 software. Extracted particles were used for iterative 2D-classification followed by the elimination of bad classes. Consolidated information for the analyzed data (micrographs, particles, numbers of classes) is presented in Table S2. Linear dimensions of the 2D-classes were measured with ImageJ (*38*). Initial 3D reconstitution was performed in EMAN2.3 (*23*) using selected classes representing different views of the particles. Initial 3D-models were imported to RELION 2.1 and used as a reference for 3D classification, followed by auto-refinement, masking, post-processing and determination of final resolution in this program. All 3D reconstructions were visualized and analyzed in UCSF Chimera (*39*). Analysis of the FACT-ΔNTD mutant was conducted using the same steps, except single particle images were collected using autopicking utility in RELION3.0 software.

### Structural analysis of the interacting domains in the compact FACT conformation

The structure of the three-domain FACT complex was constructed based on the crystal structure of FACT-nucleosome complex 2 (pdb id 6upl (*17*)). The model of unfolded FACT-nucleosome complex was built using the atomic structure of the folded complex (pdb id 6upl (*17*)), followed by rigid fitting of the domains with DNA in the map of electron density of the complex using USCF Chimera software. The structure of *Homo sapiens* NTD-Spt16 domain was downloaded from rscb.org (pdb id 5e5b (*20*)). The hydrophobic organization of interacting subunits in compact four-domains structure FACT was analyzed using Platinum web service (*40*) and UCSF Chimera (*39*). The data on hydrophobic organization of interacting subunits were used to determine the initial orientations of the domains for flexible molecular docking via HADDOCK 2.4 server (*41*). Based on the HADDOCK score and RMSD (root mean square deviation) all conformations were divided into clusters and were visually analyzed in UCSF Chimera. The contacts between subunits of FACT were evaluated using the Protein Interactions Calculator (PIC) server (*42*).

### Statistical Analysis

In spFRET measurements, the EPR profiles and contents of nucleosome subpopulations were averaged (mean±SEM) over three independent experiments. The sample sizes varied from 1600 to 8800 particles per each independent experiment.

In electron microscopy experiments, fractions of open and closed complexes were calculated as the average of three experiments.

## Supporting information

supplement images

## Acknowledgments

Authors would like to thank Dr. G. Mer for providing the mutant FACT, Dr. K. Luger for providing the 2D class averages of the FACT-nucleosome complex, Dr. A. V. Feofanov for help with spFRET analysis and E. Trifonova for critical comments on the manuscript. V.M.S., O.I.V., O.S.S. are members in Interdisciplinary Scientific and Educational School “Molecular Technologies” of the Living Systems and Synthetic Biology of Moscow Lomonosov University. Electron microscopy was performed using the Unique equipment setup “3D-EMS” in the Moscow State University.

## Funding

This work was supported by:

National Institutes of Health Grant R01 GM119398 (VMS)

National Institutes of Health Grant R01 GM064649 (TF)

RSF (19-74-30003) (MPK, electron microscopy and spFRET experiments)

RFBR (20-54-04004) (AVP, molecular modeling experiments)

## Author contributions

Conceptualization: VMS, OSS, MPK

Methodology: VMS, OSS

Investigation: OIV, ALS, AVM, AVP, MGK, MV, EYK

Visualization: OIV, ALS

Supervision: VMS, OSS

Writing—original draft: OIV, ALS, AVP, VMS, OSS

Writing—review & editing: VMS, OSS, TF

## Competing interests

Authors declare that they have no competing interests.

## Data and materials availability

The datasets generated and analyzed during the current study are available from the corresponding authors on reasonable request.

## Supplementary Materials

Please see separate pdf file.

## Notes

### Competing Interest Statement

The authors have declared no competing interest.

