## supplement images for "Mechanism of Curaxin-dependent Nucleosome Unfolding by FACT"

**This PDF file includes:**

Figs. S1 to S10
Tables S1 to S2

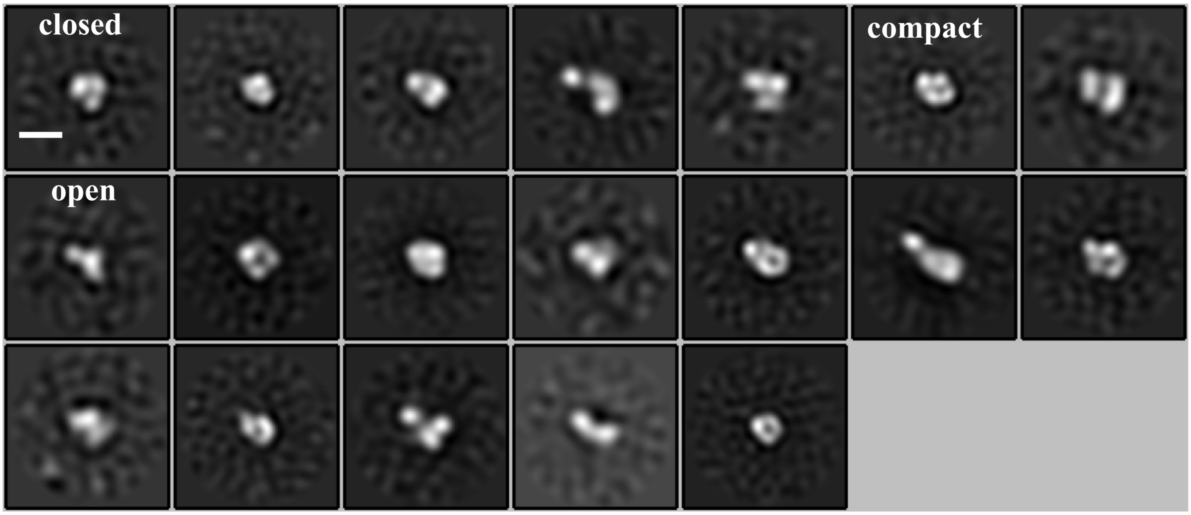


**Fig. S1. 2D class-averages of FACT complexes.**
Three distinct conformations (compact, closed and open) were identified. Bar – 10 nm.

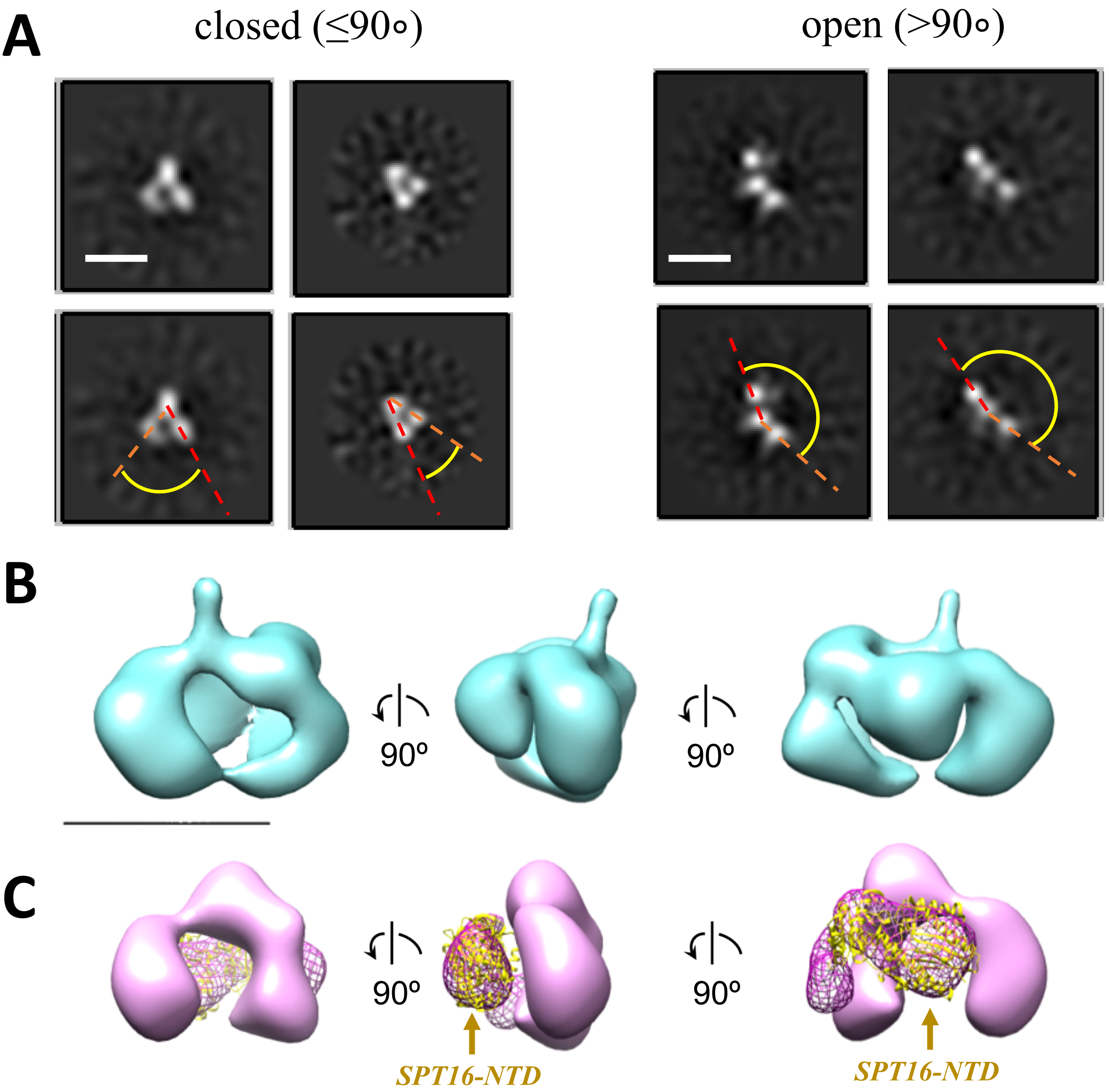


**Fig. S2. Structure of FACT-SPT16ΔNTD truncated mutant.**
**A.** Representative 2D class-averages of the FACT-ΔNTD in the closed and open conformations. Scale bars – 10 nm. **B.** 3D map of FACT in the compact conformation. **C.** 3D map of the closed conformation of FACT-ΔNTD mutant (pink, the same orientations as in part **B**) and the difference map with the intact FACT (magenta mesh). Atomic model of SPT16-NTD (yellow ribbon) was docked into the difference map with the correlation coefficient 0.92. (**C**) of in the same orientations as structure in (**B**). Bar – 10 nm.

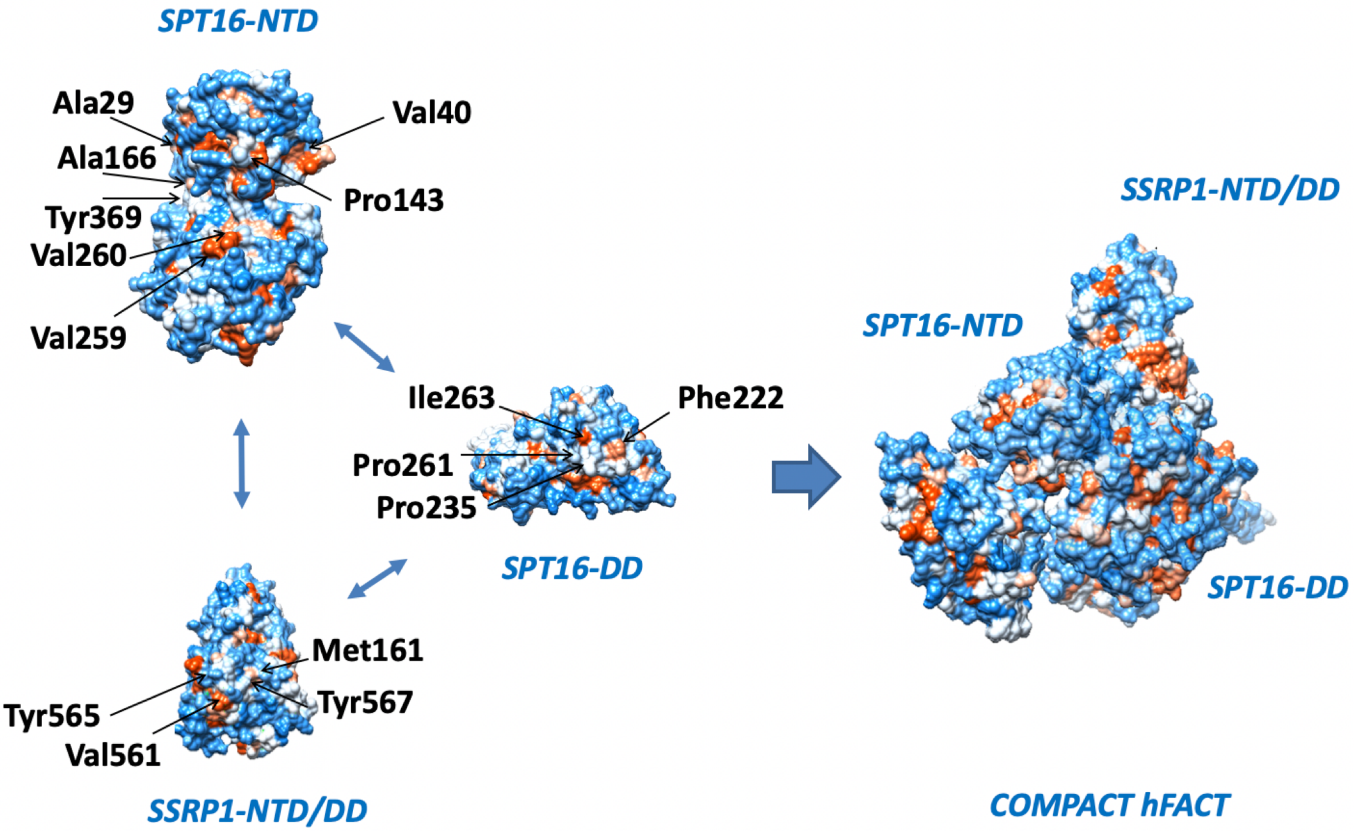


**Fig. S3. Surfaces participating in interactions between domains of human FACT subunits.**
The interacting surfaces of different FACT domains are shown on the left and the entire compact FACT structure in the compact conformation – on the right. Hydrophobic and hydrophilic regions are shown in orange and blue, respectively. The key amino acids involved in the interactions are indicated.

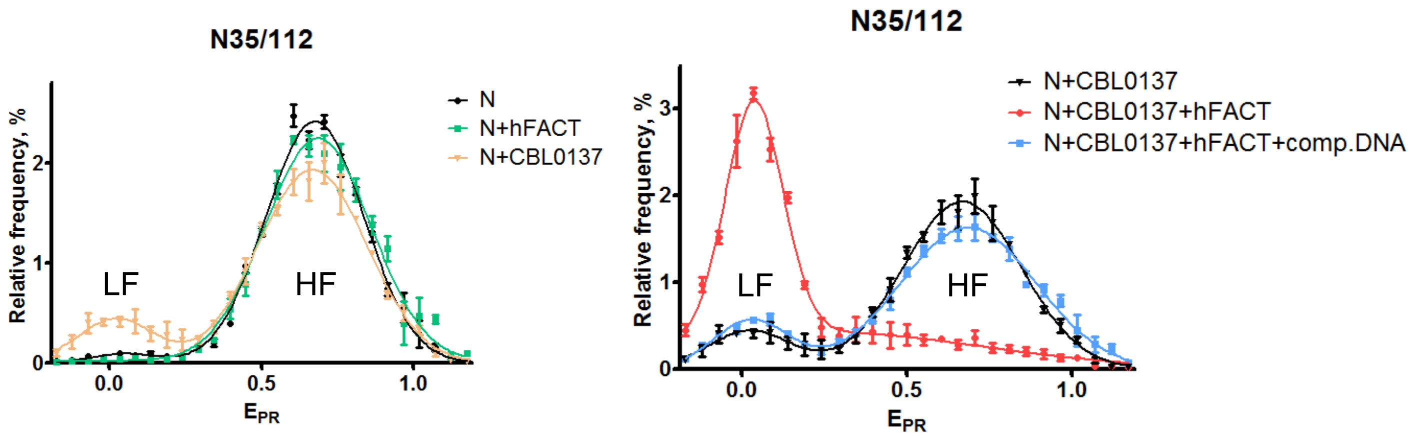


**Fig. S4. FACT and curaxin CBL0137 work synergistically and induce a large-scale, reversible nucleosome reorganization**.

Typical frequency distributions of FRET efficiencies (EPR) in the presence or absence of curaxin CBL0137, FACT and/or competitor DNA. Analysis by spFRET microscopy. The mean values of EPR peaks and the standard errors were the following: (N) – 0.057±0.063, 0.676±0.004; (N+FACT) – 0.073±0.093, 0.688±0.014; (N+CBL0137) – 0.027±0.048, 0.669±0.012; (N+CBL0137+FACT) – 0.036±0.005, 0.525±0.09; (N+CBL0137+FACT+competitor DNA) – 0.026±0.005, 0.687±0.008. Low-FRET and high-FRET peaks are indicated as HF and LF, respectively.

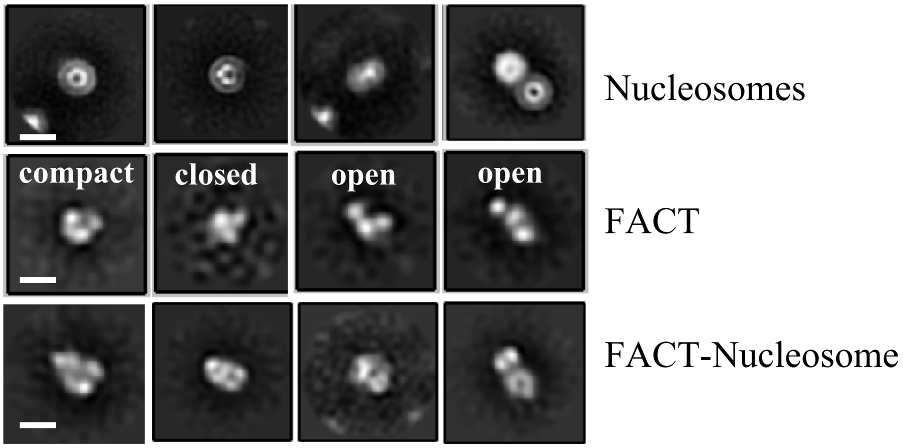


**Fig. S5. Representative 2D class-averages of complexes formed in the presence of FACT and nucleosomes without curaxin CBL0137.**

Middle raw - three distinct conformations (compact, closed and open) of FACT. Bottom raw – different views of the folded FACT-nucleosome complex. Bar – 10 nm.

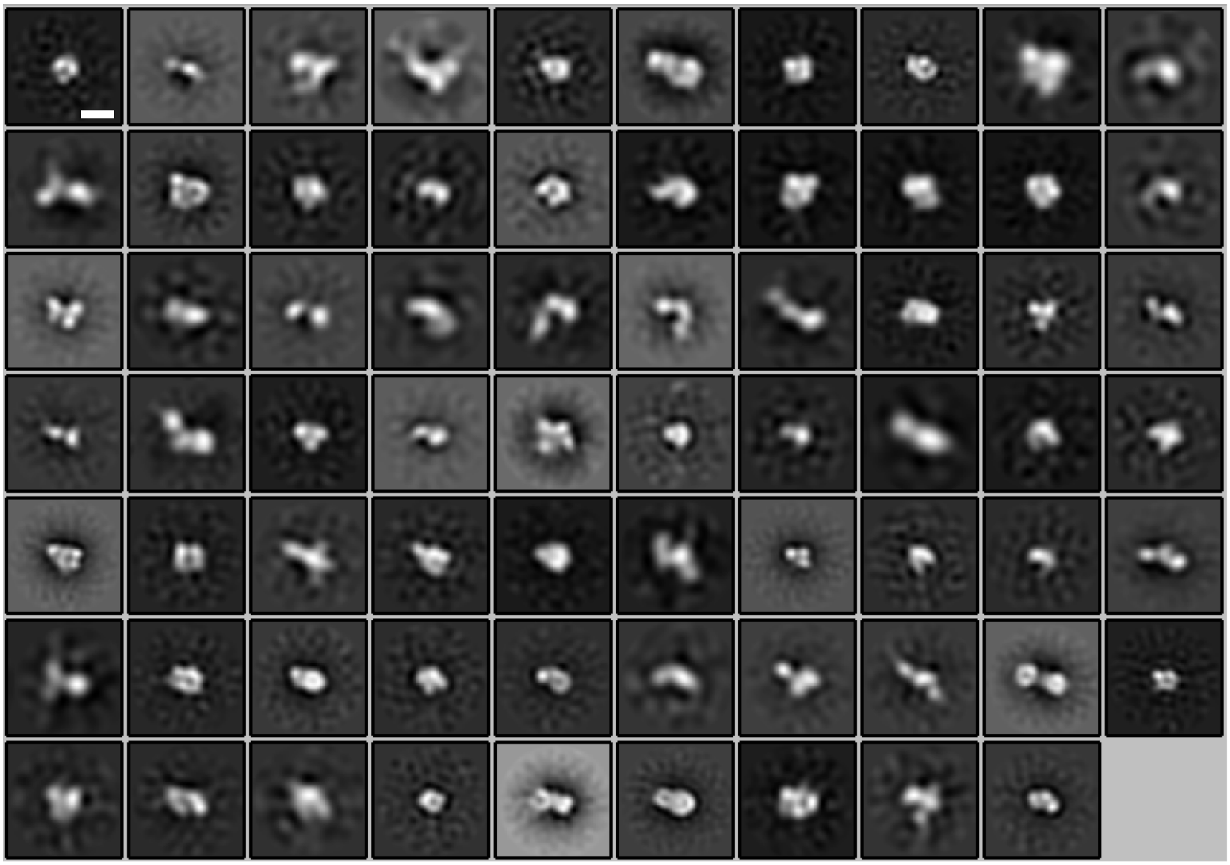

**Fig. S6. 2D class-averages of FACT complexes with nucleosomes formed in the presence of curaxin CBL0137.**

Nucleosome-free FACT, FACT-nucleosome complexes and nucleosomes are present in the sample. Bar – 10 nm.

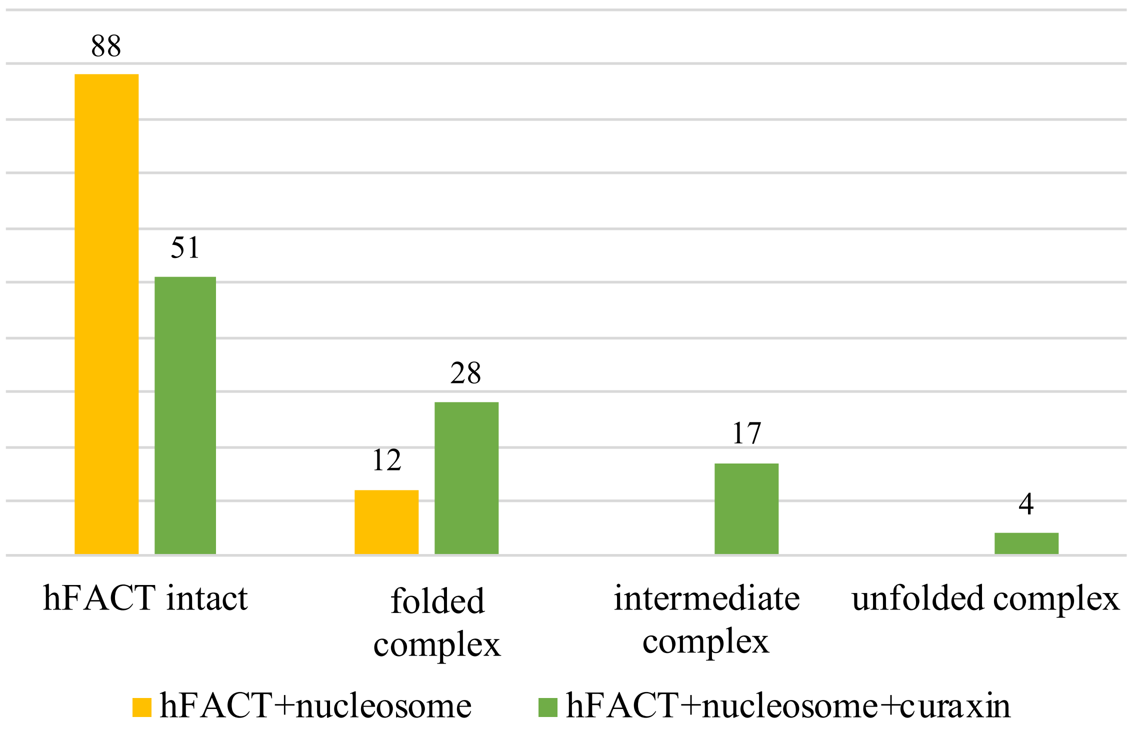


**Fig. S7. Nucleosome unwrapping by FACT in the presence of curaxin CBL0137.**
Fractions (%) of FACT-nucleosome complexes having different conformations in the absence (yellow) and in the presence (green) of CBL0137 are shown. Intact FACT is not bound to nucleosomes.

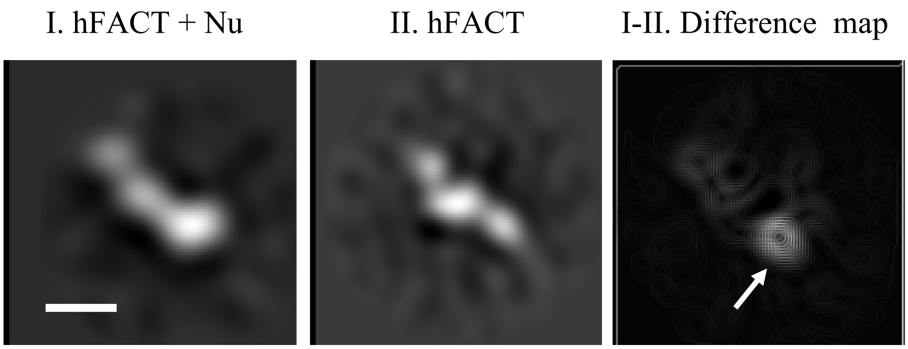


**Fig. S8. Differential map of structurally similar 2D class-averages of open FACT and FACT-nucleosome complexes**.

The differential density is indicated by arrow; it has the approximate size of the histone hexamer. Bar – 10 nm.

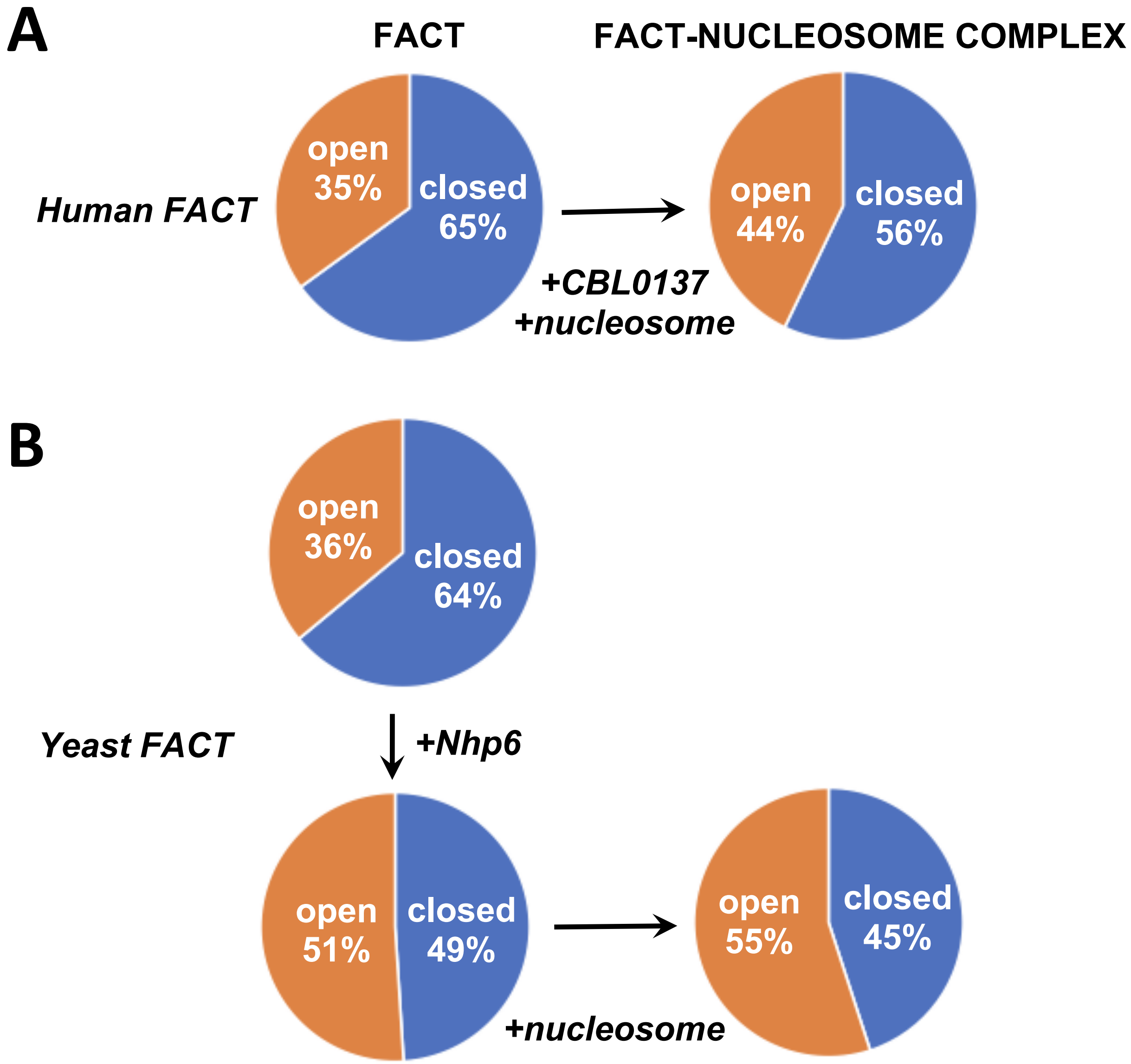


**Fig. S9. Conformations of human (A) and yeast (B) FACT in solution and in FACT-nucleosome complexes.**

To allow easier comparison between different samples, all complexes containing compact and open conformations of FACT were counted as closed and open complexes, respectively. **(A)** Human FACT-nucleosome complexes were formed in the presence of curaxin CBL0137. (**B)** Yeast FACT-nucleosome complexes were formed in the presence of DNA-binding protein Nhp6 that facilitates FACT opening, partial DNA uncoiling from histone octamer and greatly facilitates nucleosome unfolding *(7)*.

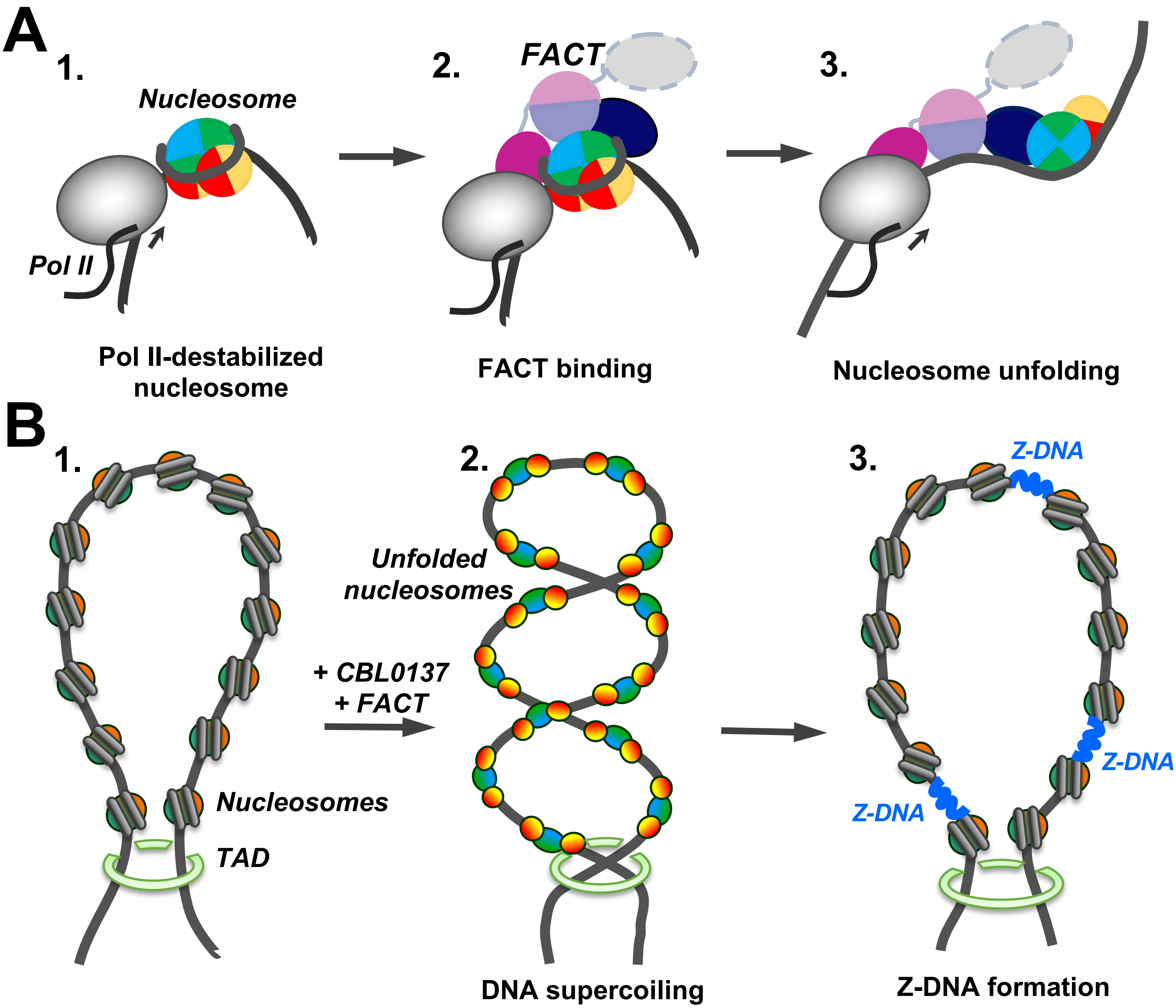


**Fig. S10. Possible functions of FACT-induced nucleosome unfolding.**
The color code as in **Fig. 5**. (**A**) FACT-dependent nucleosome unfolding during transcription by Pol II. Transcribing RNA polymerase II (Pol II) partially uncoils nucleosomal DNA from the histone octamer and opens FACT-interacting surfaces of H2A/H2B dimers (intermediate 1) *(28)*. FACT binds to the destabilized nucleosomes (intermediate 2) and induces nucleosome unfolding (intermediate 3) that could facilitate further transcription and nucleosome survival during this process. (**B**) Curaxin-dependent Z-DNA formation. As curaxins and FACT interact with nucleosomes in a chromatin region that is closed by topologically associating domains (TADs, structure 1), bulk nucleosomes are unfolded, releasing unconstrained DNA supercoiling (intermediate 2). DNA supercoiling, in turn, induces formation of Z-DNA (intermediate 3) that could serve as a trigger of necroptosis in cancer cells *(15, 30)*.

| **Sample** | **Micrographs acquired** | **Total particles number** | **Number of particles used for 3D analisys** |
|---|---|---|---|
| hFACT | 1035 | 129 197 | 67 347 |
| hFACT-(SPT16ΔNTD) | 415 | 122 548 | 33 772 |
| hFACT+nucleosome | 1092 | 216 886 | 77 630 |
| hFACT+nucleosome+CBL037 | 1965 | 151 846 | 95 634 |

**Table S1. Numbers of particles analyzed for different samples.**

| Spt16 NTD | SPT16 DD domain | SSRP1 NTD\DD domain | SSRP1 MD domain |
|---|---|---|---|
| **Hydrogen bonds** | | | |
| Asp26 | **Glu562**<br>**Gly563**<br>**Tyr565**<br>**Arg601** | | |
| Glu27<br>Asn30 | Asp564 | | |
| **Glu43** | | | Lys264<br>Arg269 |
| Glu42<br>Glu145 | | | **Lys233** |
| Asp 324 | Lys 596 | **Arg164** | |
| Lys321<br>**Lys370** | Glu597 | **Glu162**<br>**Glu149** | |
| Arg317<br>Lys321<br>**Lys370** | | **Glu145** | |
| Lys23<br>His318<br>Lys372<br>Lys373 | | **Glu149** | |
| **Glu311**<br>Asp324<br>Asp412 | | **Lys143** | |
| Asp331 | **Lys555** | | |
| Glu315 | Arg581 | **Lys143** | |
| Lys334 | Asp649 | | |
| **Hydrophobic contacts** | | | |
| Tyr170<br>**Tyr369** | | Met161 | |
| Val40<br>Pro143 | | | Phe222 |
| Pro143 | | | **Pro261**<br>Ile263 |
| Ala166 | Val561 | | |
| Val167<br>Tyr28<br>Ala29<br>**Tyr369** | **Tyr565** | | |
| Ala166<br>Val167<br>Tyr170<br>**Tyr369** | **Tyr567** | | |

**Table S2. Amino acids participating in interactions between domains of human FACT subunits.** Residues that are identical between yeast and human FACT are shown in pink.
